# Identification and Characterization of Novel Chikungunya Virus Polymerase Inhibitors

**DOI:** 10.1101/2025.09.16.676555

**Authors:** Peiqi Yin, Ryan Boyce, Sainan Wang, Michael Sirrine, Alexander Leach, Dillon Chu, Dariia Vyshenska, Zafer Sahin, Jenny Wong, Tahirah Moore, Devin Shane M. Lewis, Stephen C. Pelly, Dennis Liotta, Andres Merits, Alexander L Greninger, Richard K. Plemper, Margaret Kielian, Robert M. Cox

**Affiliations:** Department of Cell Biology, Albert Einstein College of Medicine, Bronx, NY, USA; Center for Translational Antiviral Research, Institute for Biomedical Sciences, Georgia State University, Atlanta, Georgia 30303, United States; Institute of Bioengineering, University of Tartu, Tartu, Estonia; Virology Division, Department of Laboratory Medicine and Pathology, University of Washington, Seattle, WA 98195; Department of Chemistry, Emory University, Atlanta, Georgia 30322, United States

## Abstract

Chikungunya virus (CHIKV) and other alphaviruses in the *Togaviridae* family are positive-sense RNA viruses and major human pathogens, causing millions of infections worldwide. In humans, alphaviruses such as CHIKV, Mayaro and Ross River viruses typically cause arthritogenic disease characterized by debilitating arthralgia, joint inflammation, fever, and rash. Although a vaccine was recently approved for use against CHIKV, no vaccines are licensed against other alphaviruses. No antiviral treatments are available to prevent or treat infections by any alphavirus. To address this unmet need, we used a CHIKV nanoluciferase reporter virus to develop a high-throughput screening assay for novel small-molecule inhibitors. From this campaign, we identified several unique inhibitors of CHIKV replication. Mechanistic characterization of two inhibitors revealed that both target the nsP4 RNA-dependent RNA polymerase, while susceptibility profiling pinpointed unique nsP4 mutations that specifically confer resistance. *In silico* docking analyses indicated potential binding poses of the inhibitors near the polymerase active site. Collectively, these results define multiple chemotypes for further development and highlight novel molecular targets within nsP4 for CHIKV inhibition.

**IMPORTANCE:** Chikungunya virus is a mosquito-borne pathogen that has caused millions of human infections worldwide, producing severe fever, rash, and long-lasting joint pain that can persist for months. Related viruses such as Mayaro and Ross River viruses also cause debilitating disease, yet no antiviral drugs are available to treat any infection caused by this family of viruses. In this study, we developed a high-throughput assay that allowed us to rapidly identify compounds capable of blocking chikungunya virus replication. We discovered new hit compounds that inhibit virus growth. In addition, we determined that two of the most promising hit candidates target the viral nsP4 polymerase. By identifying these novel inhibitors and characterizing both their mechanisms of action and resistance profiles, we have established the groundwork for future efforts to develop much needed therapies against chikungunya virus and related pathogens.

## INTRODUCTION

Alphaviruses are enveloped, single-stranded, positive-sense RNA viruses that infect humans worldwide (1). They are primarily transmitted by mosquitoes, with small mammals and birds acting as reservoirs in nature (2, 3). Alphaviruses are grouped into encephalitic versus arthritogenic based on their typical disease symptoms in humans. The arthritogenic alphaviruses include chikungunya virus (CHIKV), Mayaro virus (MAYV), o’nyong-nyong virus (ONNV), and Ross River virus (RRV), and generally cause disease characterized by fever, rash, joint pain, and arthralgia, which can persist for months (3-5). Severe cases can occasionally lead to neurological complications, fever, and death (6, 7). Encephalitic alphaviruses such as Venezuelan equine encephalitis virus (VEEV), Western equine encephalitis virus (WEEV), and Eastern equine encephalitis virus (EEEV) more commonly present with febrile illness and severe neurological disease, including life-threatening encephalitis (8). For EEEV, mortality rates can reach 50%, with sporadic outbreaks documented in the eastern United States (9).

CHIKV originated in sub-Saharan Africa and has recently spread across the globe, becoming endemic in many regions of Central America, the Caribbean, South America, Africa, and Southeast Asia (4, 10). The first documented case of CHIKV in the Americas occurred in 2013 on St. Martin Island, and over a million cases have been reported since (11). CHIKV imposes a significant health burden on millions of people, in particular since acute infections can progress to chronic, debilitating polyarthritis, with symptoms persisting for months or years (12). The spread of CHIKV is linked to the growing distribution of mosquito vectors, consequences of climate change, and viral mutations that enhance transmission efficiency by mosquito vectors (4, 10, 13). CHIKV is spread by both *Aedes aegypti* and *Aedes albopictus* mosquitoes and current models predict that hundreds of millions more people could be at risk of CHIKV infection by the end of the 21st century (14-16). Recent outbreaks in China, France and other previously unaffected regions underscore the rapidly expanding distribution of CHIKV (17, 18). Two vaccines, Vimkunya and Ixchiq, were recently approved in the United States (3). However, the FDA subsequently withdrew approval for the Ixchiq vaccine due to serious adverse events, including several deaths. Currently, there are no licensed antiviral therapies against CHIKV, and treatment is limited to nonsteroidal anti-inflammatory drugs (19) and supportive care.

Direct-acting small-molecule antivirals are a promising strategy for treating viral infections, and could offer a stable and rapid adjunct to vaccination in controlling CHIKV outbreaks. CHIKV and other alphaviruses infect cells by receptor-mediated endocytosis and low pH-triggered fusion in acidic endosome compartments (1). The RNA genome is thereby released into the cytoplasm, uncoated, and translated to generate the membrane-associated replication complex (20, 21). This complex is generated from a nonstructural (ns) polyprotein (P1234), which is processed by the essential nsP2 protease to produce nsP1-4 (1). Among these proteins, the nsP2 protease activity and the nsP4 RNA-dependent RNA polymerase (RdRp) are especially attractive targets for inhibition. Numerous strategies have been explored to inhibit nsP2, including peptidomimetics, cysteine-reactive “warheads,” and allosteric inhibitors (22-24). However, these efforts have yet to yield a hit candidate that has advanced to clinical use. The nsP4 RdRp activity is also a major target for small-molecule inhibitors, especially since nsP4 is highly conserved among alphaviruses (25). Several nucleoside analogs potently inhibit alphavirus replication, including favipiravir(26), ribavirin (27), β-D-N4-hydroxycytidine (NHC), the bioactive metabolite of molnupiravir (27), sofosbuvir (28), and 4′-fluorouridine (4′-FlU, EIDD-2749) (29, 30). To date, however, none have yet been licensed for clinical use. A drawback to nucleoside analog therapies is their inherent potential for toxicity through incorporation by host cell polymerases, which is especially significant when treating neonatal populations, a group that, along with immune suppressed patients and the elderly, is particularly susceptible to severe CHIKV disease. Allosteric inhibitors of nsP4 could therefore represent an important strategy for safer, more targeted antiviral therapies but to date no small-molecule allosteric inhibitors of CHIKV nsP4 have been identified.

Here we conducted high-throughput screening (HTS) using a CHIKV nanoluciferase reporter virus followed by an extensive counter-screening campaign. We identified several hit compounds with anti-CHIKV activity and demonstrated that they were also active in CHIKV replicon assays. Further studies of two hit compounds showed that they inhibited CHIKV replication in normal human dermal fibroblasts and CHIKV production in human cells. Both compounds showed inhibition against a subset of alphaviruses in trans-replicase assays. Resistance profiling revealed that mutations in nsP4 reduce viral susceptibility to these compounds. *In silico* docking suggested interactions with two previously uncharacterized sites in the viral polymerase. Collectively, these findings reveal new chemotypes targeting unique microdomains of CHIKV nsP4 and provide a foundation for the future development of small-molecule polymerase inhibitors.

## RESULTS

### HTS campaign

We developed an HTS protocol using a recombinant CHIKV (181/25 strain) that expresses nanoluciferase (CHIKV-nLuc) (Fig. 1A). After validating this assay (Fig. 1B-C), we screened a 410,000-compound library (Fig. 1D–E). The HTS campaign performed well, with mean Z’=0.5 and an average signal-to-background ratio of 32. By combining control-dependent methods (normalized percent inhibition, NPI) and control-independent methods (robust Z-score) (Fig. 1E), we identified 740 compounds for secondary orthogonal counter-screens in a 384-well format (Fig. 1F). The counter-screen criteria required EC_50_ <5 µM and CC_50_ >20 µM. Compounds showing activity against unrelated viral families, causing reporter interference, or exhibiting structural liabilities were excluded. From these 740 compounds, 14 hits, representing 10 distinct chemotypes, met all performance criteria in 384-well dose-response assays (Fig. 2). These hits were then procured for 96-well dose-response confirmation (Table 1). Three chemotypes emerged that merited further investigation. However, one chemotype, represented by GAP-1066658, was previously identified as an inhibitor of the CHIKV capping machinery (31, 32). Although we excluded this chemotype from additional testing, it validated our anti-CHIKV HTS platform. The remaining two chemotypes, GAP-1146924 and GAP-1173149, were advanced for mechanistic investigations.

**Figure 1.**
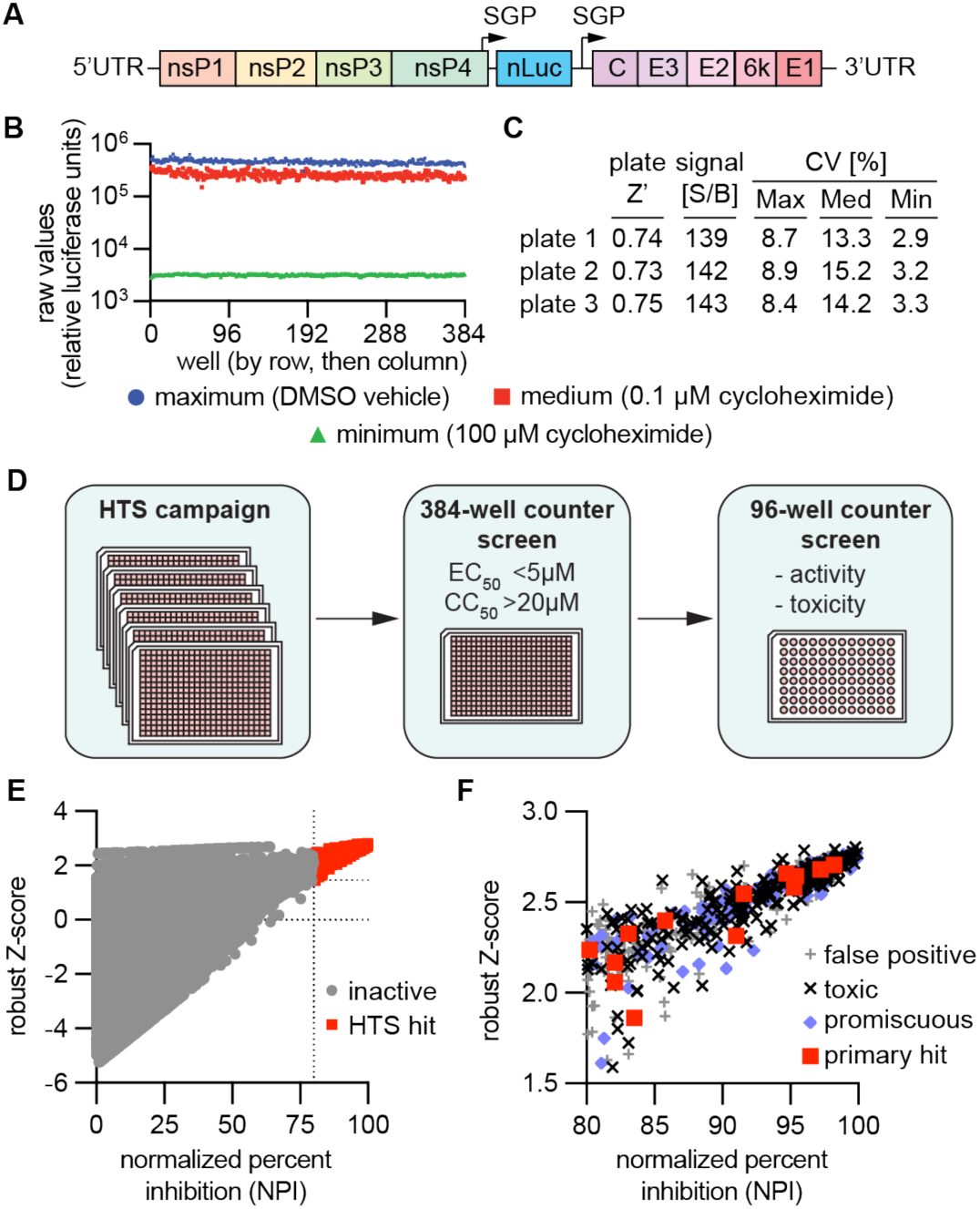
CHIKV HTS campaign. **A**) Diagram of the recombinant CHIKV reporter virus (CHIKV-nLuc) expressing nanoluciferase from the viral subgenomic promoter (SGP) and structural proteins from an additional SGP inserted between the nLuc and capsid ORFs that was used to develop a HTS assay. **B**) Validation of automated HTS protocol in 384-well format, using CHIKV-nLuc. Three 384-well assay validation plates featuring alternating columns of DMSO vehicle (0.1%; max/maximum) or the broad-spectrum host-directed inhibitor cycloheximide in intermediate (0.5 × EC_50_ (0.1 µM); med/medium) or sterilizing (10 × EC_50_ (100 µM); min/minimum) concentrations. On each validation plate, control columns were arranged in different order. **C**) Performance parameters for all three plates and statistical assessments are shown. Signal-to-background (S/B), coefficients of variation for max, med, and min signals, and Z’ scores for each reference plate are shown. **D**) Schematic of the HTS process. **E**) Compounds were initially tested in single concentration (5 µM), 384-well format. Primary hit analysis was performed using control dependent (normalized percent inhibition) and control independent (robust Z-score) methods. **F**) Primary hits were tested in 384-well dose response format for primary target activity, toxicity and activity against unrelated viruses to test for anti-CHIKV activity, toxicity, and promiscuity, respectively.

**Figure 2.**
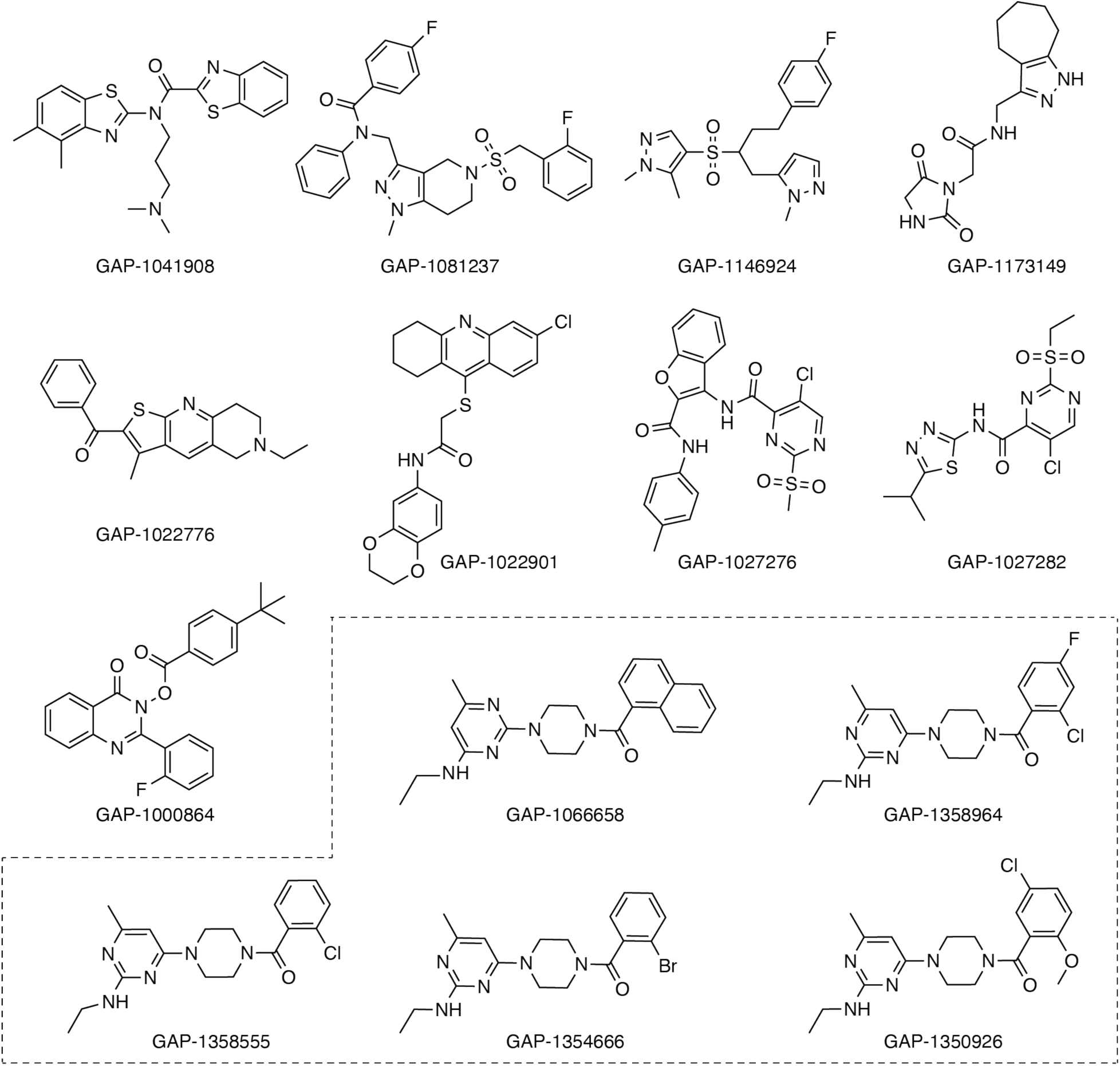
Primary screening hits. 14 compounds representing 10 unique scaffolds passed initial 384-well dose response counterscreens. These compounds were sourced from their respective vendors and selected for additional 96-well dose response assays. Compounds belonging to the anti-CHIKV chemotype previously identified by Abdelnabi et al.(31) are outlined (dashed line).

**Table 1.** Bioactivity profiles of primary hit candidates.

| Compound ID | Antiviral potency (EC<sub>50</sub> [μM]) |  |  |  |  |  |  |  |  | Toxicity (CC<sub>50</sub> [μM]) |  |
| --- | --- | --- | --- | --- | --- | --- | --- | --- | --- | --- | --- |
|  | CHIKV (U-2 OS) | CHIKV (Vero) | CHKV replicon | VEEV | SINV | HPIV3 | SARS-CoV-2 | RSV | VSV | VeroE6 | U-2 OS |
| GAP-1041908 | 13.3 | 22.2 | 32.1 | 18.8 | 18.9 | 41 | 28 | -- | 32.7 | 34.9 | 31.4 |
| GAP-1081237 | 9.1 | 10.5 | 12.4 | 10.0 | 65.7 | 64.9 | >100 | >10 | 65.2 | >100 | >100 |
| **GAP-1146924** | 24.1 | 19 | 9.2 | 35.2 | >100 | >100 | >100 | >10 | >100 | >100 | >100 |
| **GAP-1173149** | 15.6 | 18.6 | 5.3 | 52.3 | >100 | >100 | >100 | >10 | >100 | >100 | >100 |
| GAP-1022776 | 9.1 | 7.7 | n.d. | n.d. | >20 | >20 | >20 | >10 | >20 | 25.2 | 19.2 |
| GAP-1022901 | >10 | 3.76 | n.d. | n.d. | >20 | >20 | >20 | >10 | >20 | >20 | >20 |
| GAP-1027276 | ~10 | ~10 | n.d. | n.d. | >20 | >20 | >20 | >10 | >20 | >20 | 20 |
| GAP-1027282 | 1.9 | >10 | n.d. | n.d. | >20 | >20 | >20 | >10 | >20 | >20 | 16.7 |
| GAP-1000864 | ~10 | >10 | n.d. | n.d. | >20 | >20 | >20 | >10 | >20 | >20 | >20 |
| **GAP-1066658** | 3.1 | 3.2 | 0.61 | 75.8 | >100 | ~100 | ~100 | >100 | 60.9 | 88.5 | 61.5 |

Note: SINV, HPIV3, VSV, and VEEV potency was assessed on VeroE6 cells. RSV potency was assessed on Hep-2 cells. SARS-CoV-2 potency was assessed on VeroE6-TMPRSS2 cells.

### Indication spectrum and mechanistic characterization

GAP-1146924 and GAP-1173149 both showed activity that appeared specific to alphaviruses, as neither compound inhibited human parainfluenza virus 3 (HPIV3), respiratory syncytial virus (RSV), SARS-CoV-2, or vesicular stomatitis virus (VSV), and neither displayed detectable toxicity at concentrations exceeding 100 µM (Fig. 3A, Table 1). Neither inhibited the alphavirus Sindbis virus (SINV), which is relatively distant from CHIKV. Time-of-addition (ToA) studies using CHIKV-nLuc were then conducted to gain preliminary insight into which stage of viral replication is targeted by these compounds (Fig 3B). Previously characterized CHIKV polymerase inhibitors 4′-FlU and EIDD-1931 (27, 29) served as polymerase-inhibition controls, while ammonium chloride treatment was used as an entry-inhibition reference (33). The ToA profiles of GAP-1146924 and GAP-1173149 closely resembled those of 4′-FlU and EIDD-1931, suggesting these compounds act on the viral RdRp (Fig. 3B). To evaluate their potential as CHIKV replicase inhibitors, we initially tested GAP-1146924 and GAP-1173149 in dose response replicon assays. Both compounds exhibited low micromolar potency (EC_50_ of 9.2 µM and 5.3 µM for GAP-1146924 and GAP-1173149, respectively) in replicon assays, providing additional support for the hypothesis that they inhibited viral RNA synthesis.

**Figure 3.**
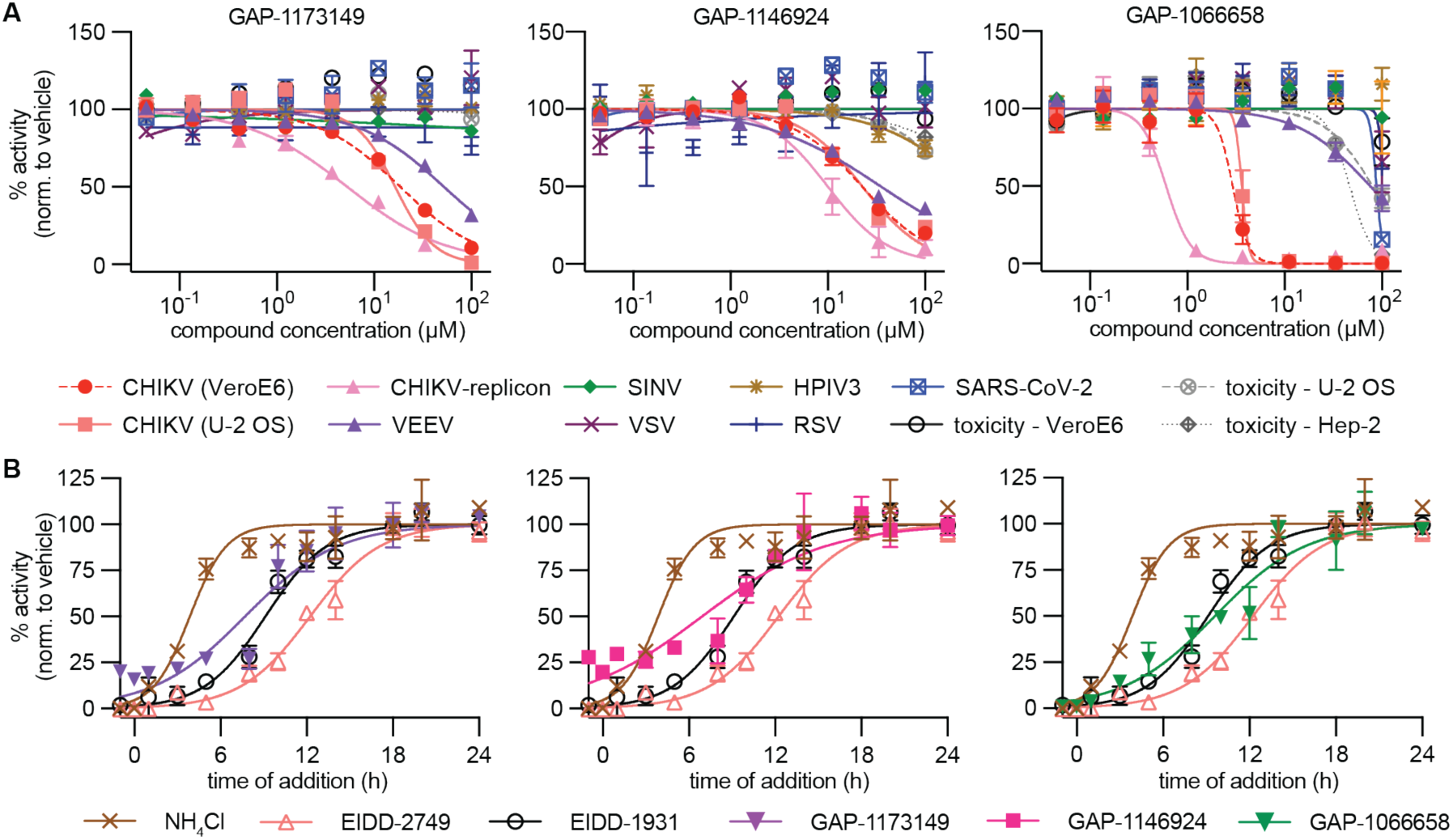
Validated Hit scaffolds. **A**) 96-well dose response testing was performed on GAP-1173149 (left), GAP-1146924 (center), GAP-1066658 (right). GAP-1173149 and GAP-1146924 displayed alphavirus-specific activity with no measurable toxicity (CC50>100 µM; 72 h incubation). **B**) Time-of-addition profiles of GAP-1173149 (purple triangles, left), GAP-1146924 (magenta squares, center), GAP-1066658 (green triangles, right). EIDD-1931 (black open circles) and 4′-FlU (pink open triangles) were used as internal references for polymerase inhibitors. Ammonium chloride was used as a reference for entry inhibition. Time-of-addition studies were performed in U-2 OS cells. Data are the mean± s.d. of three independent experiments.

Next, we assessed the antiviral efficacy of GAP-1173149 and GAP-1146924 against CHIKV infection of primary normal human dermal fibroblast (NHDF) cells, as dermal fibroblasts are a key initial target of CHIKV infection (5, 34). GAP-1173149 or GAP-1146924 inhibited nLuc production with an EC_50_ of 11.62 µM or 9.33 µM respectively (Fig. 4A and 4C), similar to the EC_50_ observed in the U-2 OS cells used for the HTS. Analysis of GAP-1173149 and GAP-1146924 cytotoxicity in NHDF cells showed that the CC_50_ for both compounds was greater than 500 µM (Fig. 4B and 4D), resulting in a high selectivity index (SI = CC_50_/EC_50_, >43 or >54 respectively).

**Figure 4.**
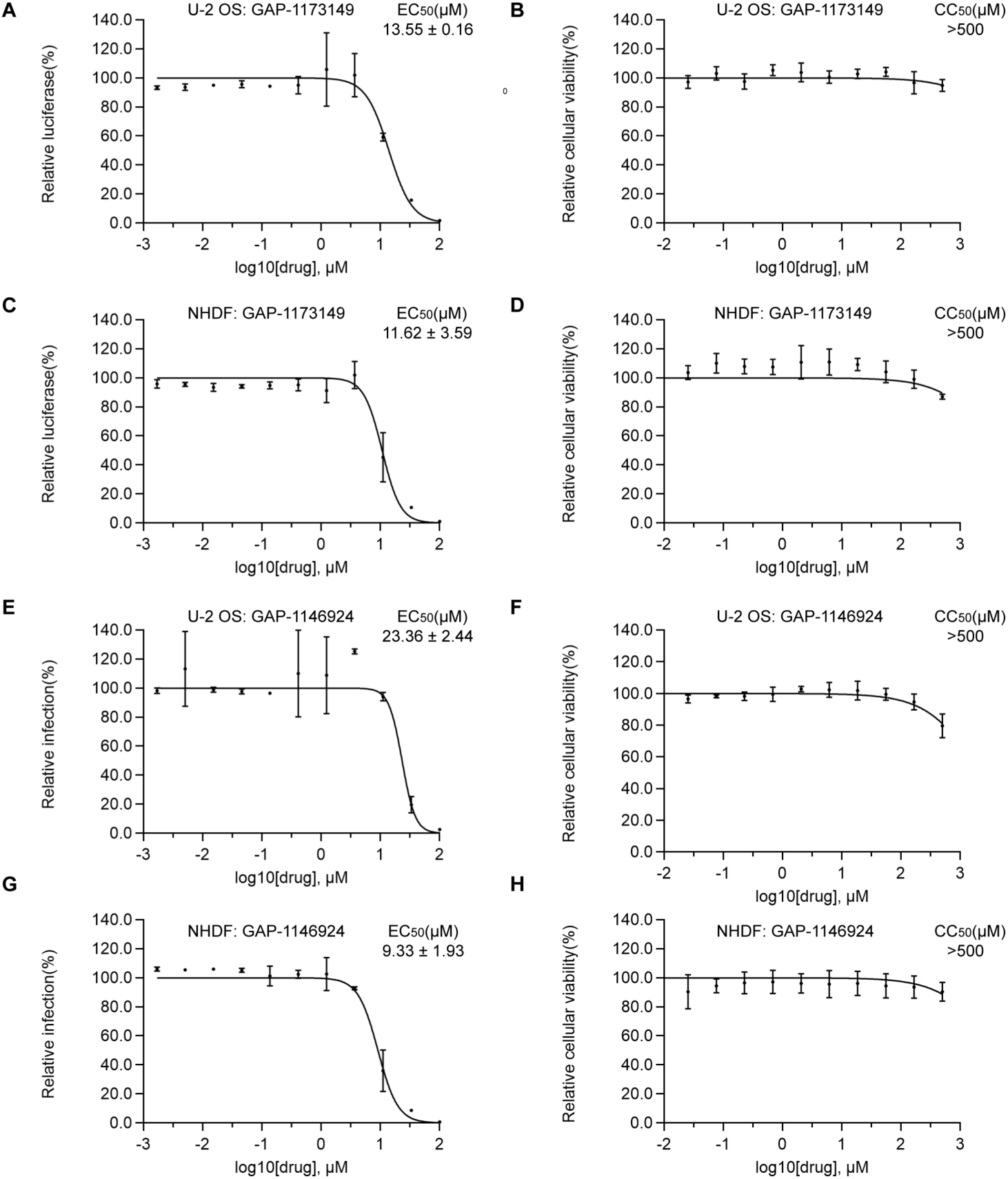
Dose-response studies of GAP-1173149 and GAP-1146924 in CHIKV-infected NHDF cells. (**A** and **C**) Effect of GAP-1173149 or GAP-1146924 on CHIKV-nLuc. NHDF cells were infected with CHIKV-nLuc at a MOI= 0.1 FFU/cell for 1 hour and then treated with the indicated concentrations of GAP-1173149 or GAP-1146924. Nanoluciferase activity was measured at 24 hpi and normalized to that of control cells. (**B** and **D**) NHDF cells were incubated with the indicated concentrations of compound for 24 hours. Cell viability was assessed using PrestoBlue and normalized to control DMSO-treated cells. Symobls in all panels represent the mean± s.d. of three independent experiments.

### Antiviral breadth of GAP-1173149 or GAP-1146924 against alphaviruses

To evaluate the broad-spectrum antiviral potential of GAP-1173149 and GAP-1146924 across the alphavirus genus, we tested their inhibitory activity against a panel of re-emerging pathogens (ONNV, RRV and MAYV), as well as the less pathogenic SINV and Semliki Forest virus (SFV). U-2 OS cells were inoculated with the indicated alphavirus at a multiplicity of infection (MOI) of 0.1 focus-forming units (FFU) per cell for 1 hour. Cells were then treated with varying concentrations of GAP-1173149 or GAP-1146924, and supernatants were collected at 24 hpi and titered. GAP-1173149 effectively suppressed the replication of ONNV and RRV, showing EC_50_ values comparable to those observed for CHIKV (Fig. 5A). In contrast, GAP-1146924 demonstrated inhibitory activity against CHIKV, ONNV, and MAYV but not RRV (Fig. 5B). To further investigate the inhibitory effects on viral RNA replication across various alphaviruses, especially high-risk encephalitic alphaviruses (belonging to biosafety level 3), we employed a trans-replicase system in which expression of the P1234 polyprotein drives replication of a reporter RNA template (Fig. 5C). In this system, 100 µM GAP-1173149 effectively inhibited RNA replication of EEEV and Barmah Forest virus (BFV). It also suppressed replication of CHIKV, ONNV and RRV, but showed no significant effect on MAYV, SINV, or SFV (Fig. 5D), consistent with the virus production results shown in Fig. 5A. At the single 100 µM concentration tested, GAP-1146924 showed some inhibition of RNA replication across all the tested alphavirus constructs including WEEV, EEEV, VEEV, and Eilat virus (EILV), an insect-specific alphavirus (Fig. 5E).

**Figure 5.**
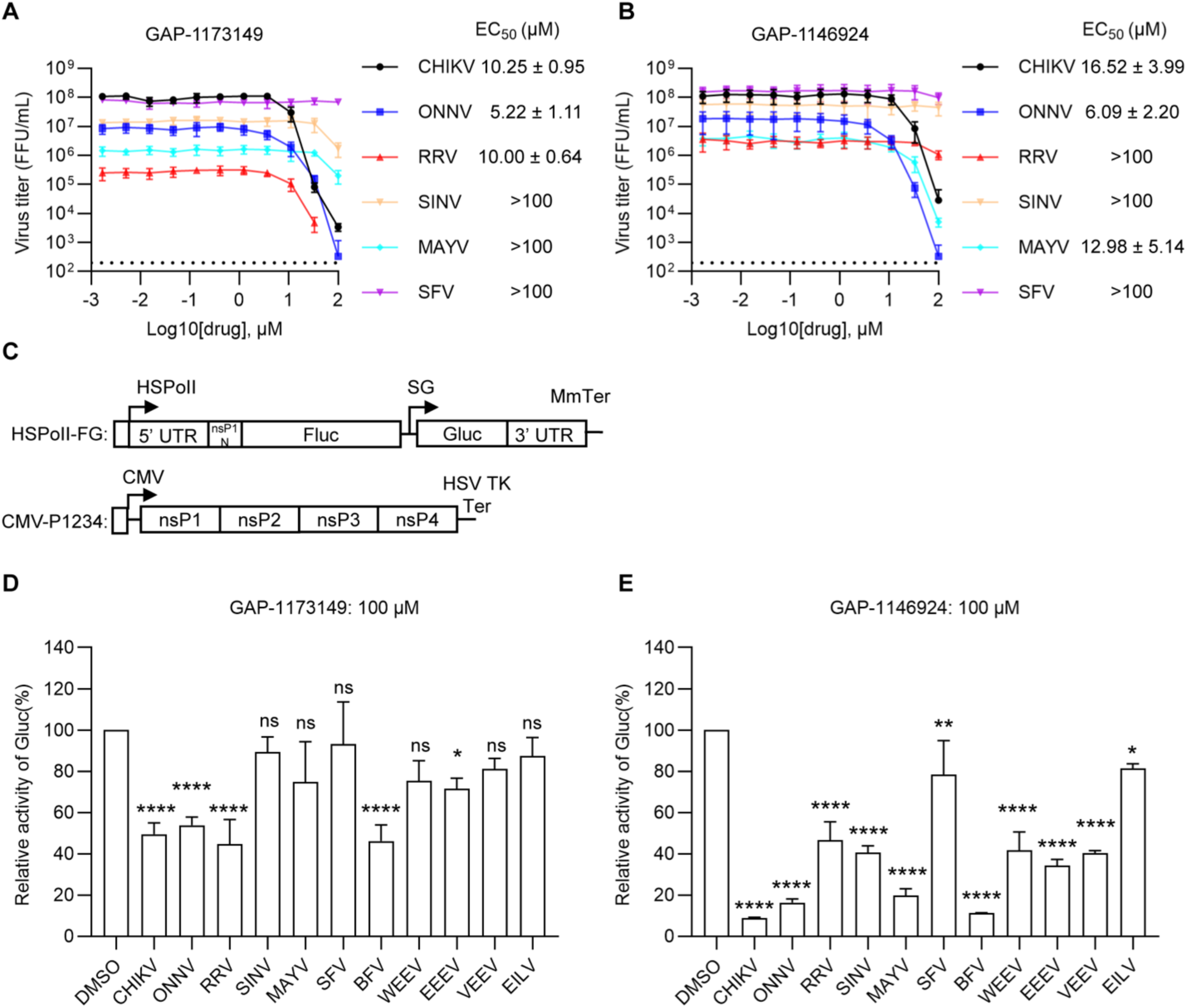
Antiviral breadth of GAP-1173149 or GAP-1146924 against alphaviruses. (**A** and **B**) Effect of GAP-1173149 (A) or GAP-1146924 (B) on the production of various alphaviruses. U-2 OS cells were inoculated with various alphaviruses and treated with the indicated concentrations of compound. Culture supernatants were collected at 24 hpi and titered by FFA on U-2 OS cells. Data are the mean± s.d. of three independent experiments. (**C**) Schematic of expression plasmids for the alphavirus *trans*-replicase assay, showing template RNA (top) and viral replicase protein expression constructs (CMV-P1234). Annotations indicate: HSPolI, a truncated human RNA polymerase I promoter (nucleotides −211 to −1); MmTer, the mouse RNA polymerase I terminator; CMV, the human cytomegalovirus immediate early promoter; HSV TK Ter, the herpes simplex virus thymidine kinase terminator. (**D** and **E**) U-2 OS cells were transfected with alphavirus *trans*-replicase constructs for 4 hours and treated with indicated concentrations of GAP-1173149 (**D**) or GAP-1146924 (**E**) for 20 hours. Gluc activity was determined and normalized to that of DMSO-treated controls. Data are the mean± s.d. of three independent experiments. Statistical significance was determined by one-way ANOVA with Dunnett’s test compared to DMSO control. \**P* < 0.05, \*\**P* < 0.01, \*\*\*\**P* < 0.0001.

### GAP-1173149 or GAP-1146924 selects for mutations in the CHIKV RdRp

To define the resistance profile and identify the viral targets of GAP-1173149 and GAP-1146924, we serially passaged CHIKV in U-2 OS cells in the presence of low concentrations of compound (Fig. 6A, 6B). Cells were inoculated at an MOI of 1 FFU/cell and cultured for 16 h in GAP-1173149, GAP-1146924 or DMSO (three independent lineages for each condition). Viral titers were quantitated after the first passage (P1), and cells were subsequently infected using the same MOI for the remaining rounds of selection. In the GAP-1173149 selected samples, virus titers at P1 were reduced by 1 log compared to DMSO-treated controls. All three GAP-1173149 lineages exhibited resistance by P4. To further select for resistance, two additional passages were performed at an increased concentration of GAP-1173149, generating P6 virus stocks (concentrations as specified in Fig. 6B). For GAP-1146924, two out of three lineages exhibited resistance to the higher concentration of drug by P5 and maintained resistance by P6. Lineage #2 of GAP-1146924 displayed some resistance by P3 but was inhibited when exposed to increased drug concentration in P4-6. RNA was extracted from all the P6 stocks, reverse transcribed, and subjected to whole-genome sequencing (WGS) using Illumina deep sequencing technology.

**Figure 6.**
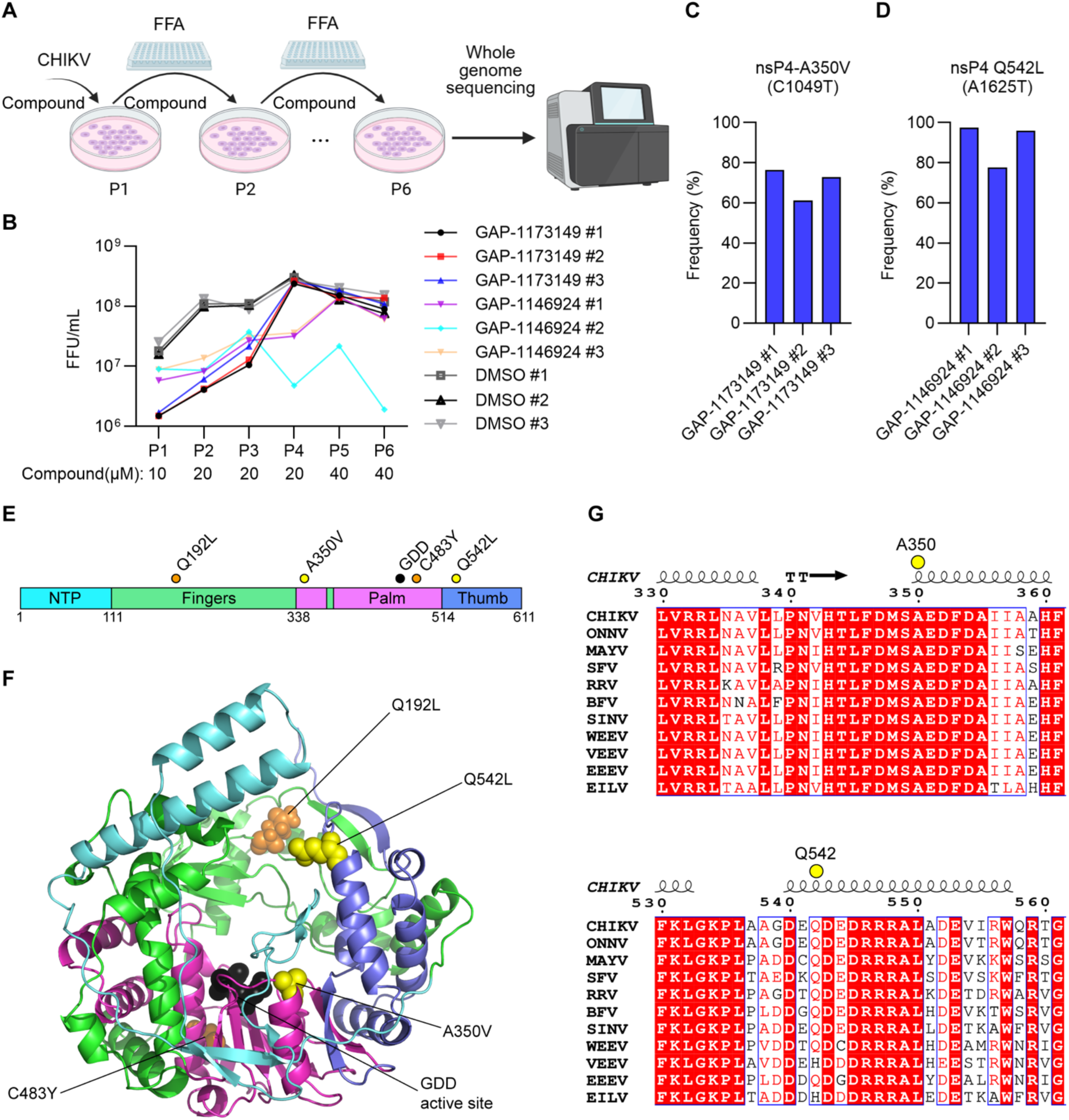
GAP-1173149 or GAP-1146924 selects for mutations in the CHIKV RdRp. **(A**) Schematic overview of the in vitro selection workflow for CHIKV under compound pressure. Figure created in BioRender. Yin, P. (2025) https://BioRender.com/ifpcoki. (**B**) Viral replication dynamics across three independent CHIKV lineages subjected to serial passaging in U-2 OS cells under indicating concentrations of GAP-1173149 or GAP-1146924 or DMSO control. Viral titers were quantified by FFA after each passage. (**C**) Mutation frequencies in viral populations from three GAP-1173149 treated lineages after passage 6, as determined by WGS. This mutation was not observed in virus populations treated with GAP-1146924 or DMSO. (**D**) Mutation frequencies in three GAP-1146924 treated viral populations after passage 6, as determined by WGS. These mutations were not detected in GAP-1173149 or DMSO treated virus populations. (**E**) Schematic of the CHIKV nsP4 polymerase. Locations of resistance mutations identified with GAP-1173149 (A350V) and GAP-1146924 (Q542L) are shown as yellow circles. Resistance mutations previously identified against 4′-FlU are shown as orange circles. The GDD active site is denoted with black circles. (**F**) Locations of antiviral resistance mutations in a 3D-model of the CHIKV nsP4 polymerase. Mutations identified with GAP-1173149 (A350V) and GAP-1146924 (Q542L) are shown as yellow spheres. Resistance mutations against 4′-FlU are shown as orange spheres. The GDD active site is shown as black spheres. The NTP, fingers, palm, and thumb domains are colored cyan, green, magenta and blue, respectively (**E** and **F**). (**G**) Alignment of alphavirus nsP4 sequences around residues A350 or Q542. These residues are indicated by yellow dots. TT stands for tight turn. Swiss-Prot accession numbers: CHIKV: A4L7I2, ONNV: P13886, MAYV: Q8QZ73, SFV: P08411, RRV: P13887, BFV: P87515, SINV: P03317, WEEV: P13896, VEEV: P36328, EEEV: Q306W6. GenBank accession number of EILV: BFZ80397.

Mutations in nsP4 were detected at high frequencies in P6 viral stocks selected with GAP-1173149 or GAP-1146924 (Table 2). All three GAP-1173149 stocks harbored an nsP4-A350V mutation at frequencies ranging from 61% to 76.3% (Fig. 6C). Lineages 1 and 3 of GAP-1146924 P6 stocks contained an nsP2-Q542L mutation at high frequencies (97.5% or 95.9%) (Fig. 6D). Lineage 2 contained both nsP4-Q542L (77.5%) and nsP4-Q192P (23.6%), with a combined allele frequency about 100% (Table 2). No mutations were identified at frequencies exceeding 10% in the DMSO-treated control lineages 1-3. The nsP4-A350V mutation is located within the palm domain and the nsP4-Q542L is within the thumb domain of nsP4 (Fig. 6E). These mutations are distinct from the previously identified Q192L and C483Y mutations that confer resistance to the nucleoside analogue 4′-FlU (Fig. 6E). In the CHIKV nsP4 polymerase structure, residue A350 is positioned adjacent to the conserved GDD active site (Fig. 6F). Interestingly, in the nsP4 structure Q542 is in close proximity to Q192, although these two residues are located in different domains. A350 is highly conserved across the alphavirus genus, whereas Q542 corresponds to a glutamate (E) in VEEV (Fig. 6G).

**Table 2.** Mutations with frequency >20% in selected CHIKV lineages.

| **Sample** | **Frequency (%)** | **Change type** | **AA change** | **Nucleotide change** |
| --- | --- | --- | --- | --- |
| GAP-1173149<br>lineage #1 | 73.77 | synonymous | NA | nsP3: A372C |
|  | 76.32 | nonsynonymous | nsP4: A350V | nsP4: C1049T |
| GAP-1173149<br>lineage #2 | 59.61 | synonymous | NA | nsP3: G1422T |
|  | 61.05 | nonsynonymous | nsP4: A350V | nsP4: C1049T |
| GAP-1173149<br>lineage #3 | 72.91 | nonsynonymous | nsP4: A350V | nsP4: C1049T |
| GAP-1146924<br>lineage #1 | 97.51 | nonsynonymous | nsP4: Q542L | nsP4: A1625T |
| GAP-1146924<br>lineage #2 | 23.56 | nonsynonymous | nsP4: Q192P | nsP4: A575C |
|  | 77.51 | nonsynonymous | nsP4: Q542L | nsP4: A1625T |
| GAP-1146924<br>lineage #3 | 95.9 | nonsynonymous | nsP4: Q542L | nsP4: A1625T |

### The nsP4 mutations specifically confer resistance to GAP-1173149 and GAP-1146924

To elucidate the role of nsP4-A350V and nsP4-Q542L, we introduced these mutations individually into the CHIKV 181/25 infectious clone, generated virus stocks, and assessed their susceptibility to GAP-1173149 or GAP-1146924. The nsP4-A350V substitution exhibited resistance to GAP-1173149, maintaining viral replication even at a concentration of 100 µM (Fig. 7A). Similarly, the nsP4-Q542L substitution showed resistance to 100 µM GAP-1146924 (Fig. 7B). Thus, each of these mutations results in appreciable CHIKV resistance. CHIKV carrying the Q192L or C483Y mutations that confer resistance to 4′-FlU (30) were not resistant to either GAP-1173149 or GAP-1146924, confirming lack of cross-resistance (Fig. 7A and B). In contrast to 4′-FlU, which functions as a nucleoside analog and viral RNA chain terminator, these findings suggest that GAP-1173149 and GAP-1146924 inhibit nsP4 by distinct mechanisms. This is in keeping with the structures of these compounds, which do not suggest that either could act as a nucleoside analogue.

**Figure 7.**
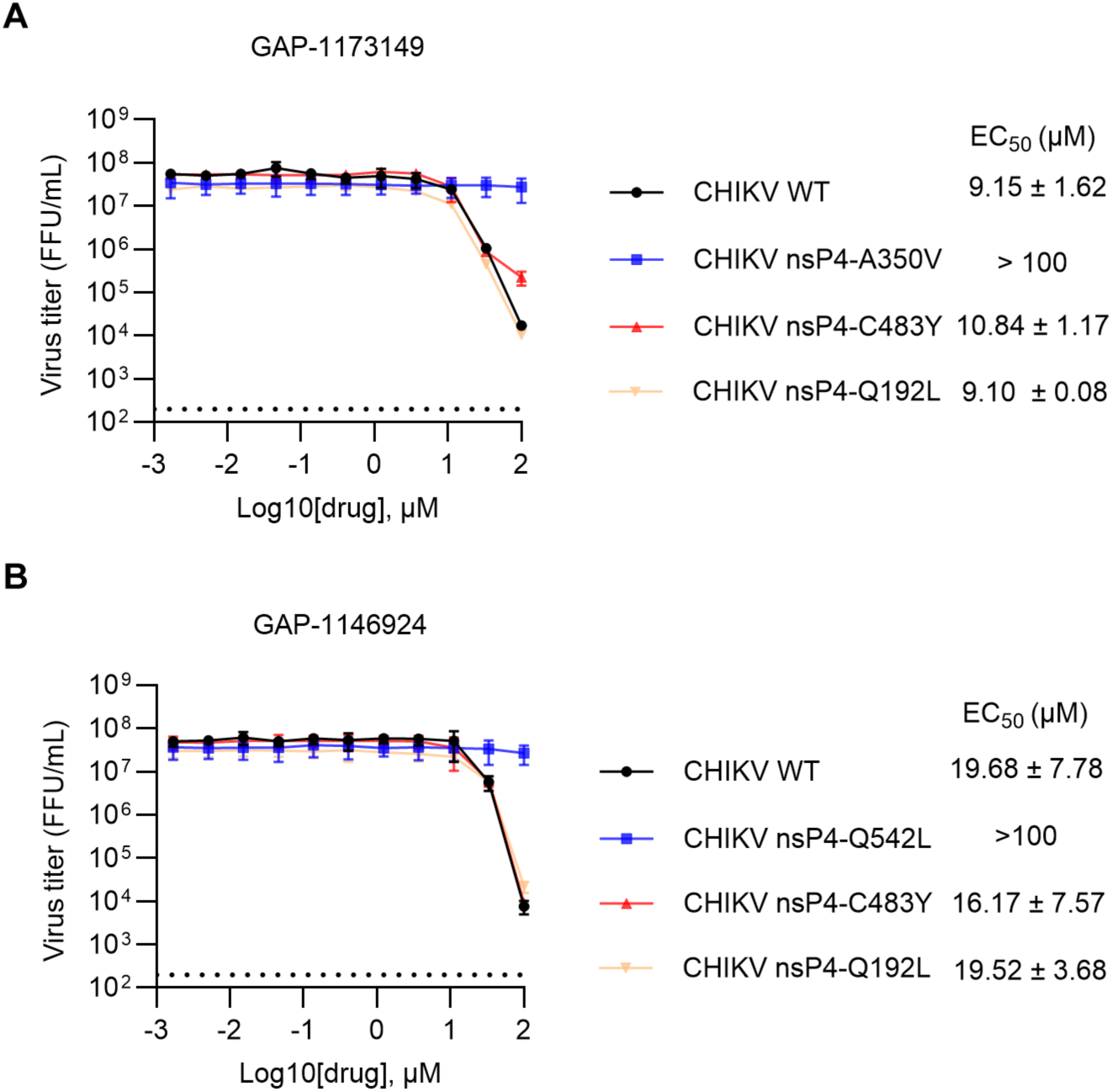
NsP4-A350V or nsP4-Q542L confers resistance to GAP-1173149 or GAP-1146924 respectively. (**A** and **B**) Targeted mutations were introduced into the CHIKV 181/25 genome, and the resulting mutant virus stocks were assessed for sensitivity to GAP-1173149 (**A**) or GAP-1146924 (**B**). Viral production was measured following treatment with the indicated concentrations, as described in Fig. 5A. Data are the mean ± s.d. from three independent experiments.

Based on the locations of mutations identified in the susceptibility profiling studies, we performed *in silico* docking using a homology model of the CHIKV nsP4 polymerase generated from the SINV nsP4 structure (PDB ID: 7VW5) and AlphaFold 3 (35). Residue A350 was selected as the docking target for GAP-1173149, and Q542 for GAP-1146924. The top-scoring docking poses for each compound are shown in Figure 8. For GAP-1173149, the pose places the compound near the GDD active site, forming a hydrogen bond between E553 and the amide group of the hydantoin ring. In this orientation, the central portion of the compound is solvent-exposed (Fig. 8G), suggesting a starting point for synthetic modifications. In contrast, the top-scoring pose for GAP-1146924 positions the compound on the opposite side of the GDD active site, near the previously reported Q192L mutation that confers 4′-FlU resistance (30). Here, a large portion of the compound is solvent-exposed, and a single hydrogen bond is observed between R593 and the sulfonyl group (Fig. 8H).

**Figure 8.**
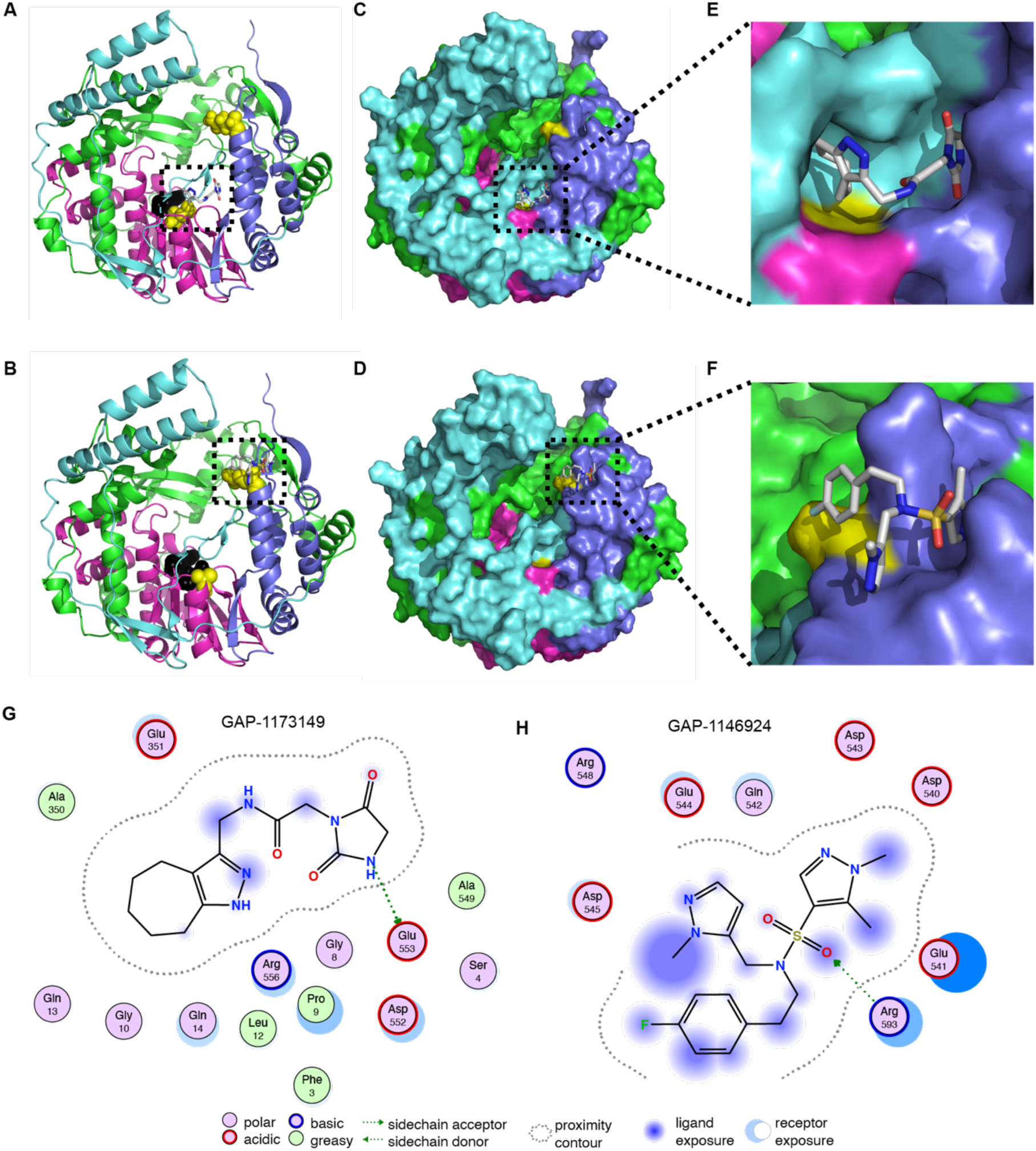
*In silico* docking. **A-B**) Location of the top-scoring docking poses for GAP-1173149 (A) and GAP-1146924 (B) are shown (dashed outline). **C-D**) Location of the docking poses within a surface representation of the CHIKV nsP4 polymerase. **E-F**) Closeup of the *in silico* docking poses for GAP-1173149 (E) and GAP-1146924 (F). Resistance mutations are shown as yellow spheres (A350V for A, C, and E; Q542L for B, D, and F). The GDD active site is shown as black spheres (A,B). The NTP, fingers, palm, and thumb domains are colored cyan, green, magenta and blue, respectively. **G**) GAP-1173149 docks within a pocket adjacent to the GDD active site. For GAP-1173149, predicted are hydrogen bond interactions between E553 and the amide group of the hydantoin ring of GAP-1173149. **H**) For GAP-1146924, the compound docks opposite of the GDD active site with large portions of the molecule being solvent exposed. Hydrogen bonds are predicted to form between R593 and an oxygen of the sulfonyl group.

## DISCUSSION

Targeting the CHIKV RdRp with small-molecule inhibitors represents a promising antiviral strategy. CHIKV nsP4 is essential for viral genome replication and transcription, making it an attractive, and highly conserved target. By inhibiting activity of the viral polymerase, replication can be halted early in the viral life cycle, shutting down viral RNA synthesis, replication, and, consequently, disease progression. In addition, the polymerase tends to be relatively well conserved among related viral species, allowing for cross-reactivity against other alphaviruses, thus enabling broad-spectrum coverage. Previous experience with other RNA virus families has demonstrated that direct-acting polymerase inhibitors can be highly effective while retaining acceptable safety profiles. For instance, polymerase inhibitors have significantly improved clinical outcomes for HIV, hepatitis C virus and other RNA viruses (36-41). Although a CHIKV vaccine is available, small molecule inhibitors can provide an additional measure to supplement protection in areas where vaccination coverage is low. This is particularly important in outbreak situations, when immediate therapeutic intervention is needed to control the spread of disease.

Here we have identified and validated two hit chemotypes, GAP-1173149 and GAP-1146924, that inhibit CHIKV in live virus assays in both immortalized human cell lines and primary human cells. Both hit candidates exhibited broad activity against multiple species within the *Alphavirus* genus. Initial mechanistic characterization suggested that the viral replication machinery was the target of inhibition. Both GAP-1173149 and GAP-1146924 inhibited RNA replication in trans-replicase assays of CHIKV, ONNV, MAYV, BFV, and VEEV. GAP-1146924 also inhibited RNA replication in trans-replicase assays of the encephalitic alphaviruses. To map the viral protein target and assess resistance barriers, we passaged CHIKV in the presence of increasing concentrations of either GAP-1173149 or GAP-1146924. We identified mutations within the CHIKV nsP4 protein and confirmed that they confer reduced sensitivity to the inhibitor. Together our data indicate that these two compounds target distinct and highly conserved microdomains within the CHIKV nsP4 polymerase. Their locations and chemotypes suggest that they act as allosteric inhibitors of nsP4.

A primary concern for the clinical performance of small molecule antivirals is the emergence of drug resistance. However, characterizing resistance mechanisms early in drug development establishes a robust foundation to inform medicinal chemistry, with the objective to potentially boost both potency and barriers to resistance. By systematically characterizing polymerase-drug interactions, lead scaffolds could potentially be refined proactively to address viral escape mutants while preserving antiviral activity. In addition, combinatorial use of inhibitors can greatly decrease the likelihood of generation of resistance mutants within the time frame of acute CHIKV infection.

In conclusion, we have identified much-needed novel small molecule inhibitors of CHIKV RNA synthesis that target conserved microdomains within the alphavirus RdRp. These two chemotypes represent a starting point for additional development to increase potency and broad-spectrum activity. Continued efforts to identify and optimize these small molecule hit candidates will advance not only our understanding of viral replication mechanisms but also the development of novel treatment options for CHIKV and related emerging alphaviruses.

## MATERIALS AND METHODS

### Cells

Human osteosarcoma (U-2 OS; ATCC HTB-96) cells were maintained at 37°C and 5% CO2 in McCoy’s media supplemented with 10% fetal bovine serum (FBS). Human carcinoma (HEp-2; ATCC CCL-23), baby hamster kidney cells (BHK-21; ATCC CCL-10), African green monkey kidney epithelial cells (VeroE6; CRL-1586; ATCC), and VeroE6 (CRL-1586; ATCC) stably expressing human TMPRSS2 (VeroE6-TMPRSS2; BBS Bioscience 78081) were cultured in Dulbecco’s modified Eagle’s medium (DMEM) supplemented with 7.5% FBS. Normal human dermal fibroblasts (NHDF; ATCC PCS-201–012) were cultured in DMEM containing 10% FBS. HEp-2 cells are listed in the International Cell Line Authentication Committee database of commonly misidentified cell lines. The use of these cells is necessary due to their permissiveness for RSV. All immortalized cell lines are routinely checked for microbial contamination and Mycoplasma. All cells were maintained at 37 °C and 5% CO_2_.

### Molecular biology

VEEV-TC83-nLuc was derived from VEEV-TC83 by adding a NanoLuc reporter cassette under the control of an additional subgenomic promoter. The infectious cDNA clone of the CHIKV-nLuc reporter virus (181/25 strain) was generated as previously described (29). To introduce specific mutations into the infectious cDNA clones of CHIKV, site-directed mutagenesis was performed using Gibson assembly. DNA fragments containing the nsP4-A350V mutation were generated by polymerase chain reaction (PCR) amplification with following primers: BSAB1-F: CCAAATCACCGACGAGTATGATGCATATCTAGACATGGTG, nsP4-A350V-F: GCATACACTATTTGACATGTCTGTCGAGGACTTCGATGCCATTATAG, nsP4-A350V-R: CTATAATGGCATCGAAGTCCTCGACAGACATGTCAAATAGTGTATGC, and Swa1-R2: GCTAACGGTTTGCCCAATTTAAATAACCTTTTTAGCGGG. The resulting PCR products were assembled into linearized CHIKV infectious clones previously digested with BsaBI and SwaI, using the Gibson Assembly Master Mix (New England Biolabs, #E2611) following the manufacturer’s protocol. The DNA fragments containing nsP4-Q542L mutation were generated using the following primers (5’→3’): NSP4-Q542L-Swa1-F: CCCGCTAAAAAGGTTATTTAAATTGGGCAAACCGTTAGCGGCAGGTGACGAACTA GATGA and SacI-R: GGTGAACCGGCCTCCTGAGTACTGTACTGCTCCGTGGTGCC. The PCR products were assembled into linearized CHIKV infectious plasmids previously digested with SwaI and SacI, using Gibson assembly as described above.

### Viruses

VEEV-TC83-nLuc virus was rescued by transfecting (Lipofectamine MessengerMAX Transfection; ThermoFisher Scientific) BHK-21 cells with *in vitro* transcribed RNA (mMESSAGE mMACHINE SP6, ThermoFisher Scientific) in accordance with the manufacturer’s protocol. Virus was harvested after 24 hours and titers were determined using TCID50 titration. VEEV-nLuc stocks were authenticated by whole genome sequencing. HPIV3 nLuc (42), VSV-nLuc(42) and RSV-A2 L19F-Firefly (43) were amplified on VeroE6 (HPIV3, VSV) and HEp-2 (RSV) cells (MOI = 0.01 TCID50 units per cell). HPIV3, VSV, and RSV stocks were purified on a 60% sucrose cushion. The identity of rescued viruses was confirmed through whole genome sequencing. Titers of HPIV3, VSV or RSV virus stocks were determined by TCID50 titration on Vero-E6 (HPIV3, VSV) or HEp-2 (RSV) cells, respectively. SARS-CoV-2 nLuc stocks were propagated on VeroE6-TMPRSS2 cells and titered by TCID50 assay on Vero-E6-TMPRSS2 cells.

Virus stocks used in the in vitro assays were generated from the following infectious cDNA clones: CHIKV 181/25 (pSinRep5-181/25ic (44), kindly provided by Dr. Terence S. Dermody), the CHIKV 181/25 mutants described above, MAYV (MAYV-CH-mKATE), RRV (RRV-T48-ZsGreen), and ONNV (ONNV-ZsGreen) (29), SFV (pSP6-SFV4)(45), and SINV-GLuc (dsTE12Q (46), kindly provided by Dr. Beth Levine). Infectious cDNA clones were linearized with the appropriate restriction enzymes, purified, and subsequently used as templates for *in vitro* transcription. Capped RNA transcripts were synthesized using the SP6 mMessage mMachine kit (Life Technologies) in accordance with the manufacturer’s instructions. The resulting RNA was electroporated into BHK-21 cells to initiate virus recovery. Electroporated cells were incubated at 37°C for 24 hours. Virus-containing supernatants were then harvested and clarified by centrifugation at 3,000 rpm for 20 minutes at 4°C using a Thermo Scientific TX-1000 rotor. Viral titers of the resulting stocks were determined by focus-forming assay (FFA) in U-2 OS cells.

### HTS and hit identification

A total of 410,000 compounds were screened for inhibitory activity against CHIKV-nLuc (MOI = 0.3 TCID50 units per cell). HTS was carried out similar to previous reports (47) (42) with a compound concentration of 5 µM and automated reading of plates 20 h after infection. Raw data were imported into the Chemical and Biological information System (ChemInnovation Software, Inc.; CBIS) IT environment and analyzed using as described (47) (42). Hit candidates were defined as compounds with anti-CHIKV activity of >5.3 × s.d. in control-dependent normalized percent inhibition and >1.45 × s.d. in control-independent mean robust Z score calculation.

### Counterscreens

For counterscreens, compound was added in threefold dilutions (20-0.009 μM or 100-0.046 µM) to 96-well plates seeded with U-2 OS, Hep2, or Vero-E6 cells (1.1 × 10^4^ per well), followed by infection with CHIKV-nLuc, VEEV-nLuc, HPIV3-nLuc, SARS-CoV-2-nLuc, VSV-nLuc, SINV-Gluc, or RSV A2-L19F reporter viruses (MOI = 0.2 TCID50 units per cell) and automated plate reading 24-36 h after infection. Cytotoxic concentrations were determined in identical serial dilution plates after exposure of uninfected cells for 72 h to the compounds. Cytotoxicity was assessed by the addition of PrestoBlue substrate (Invitrogen) to quantify cell metabolic activity, as described (42, 47). Four-parameter variable slope regression modeling was used to determine EC_50_ and CC_50_ concentrations. To determine potential structural liabilities and compound promiscuity, hit candidates were processed *in silico* through PAINS filters (48) and aggregation predictions using Badapple2 (49).

### Time-of-addition studies

Vero-E6 or U-2-OS cells were inoculated (MOI = 0.4 TCID50 units per cell) with CHIKV-nLuc. At the specified time points relative to infection, the indicated compounds were added to the culture media at a final concentration of 20 µM. Ammonium chloride was used at a concentration of 20 mM in cell culture media. Reporter gene expression was measured at 24 h after infection.

### Focus forming assay

To quantify virus titers in vitro assays, U-2 OS cells seeded at 15000 per well in 96-well plates were inoculated with 10-fold serial dilutions of virus prepared in Med-A (minimum essential medium supplemented with 0.2% BSA and 10 mM HEPES) for 2 hours. After infection, the cells were overlaid with 1% carboxymethylcellulose in modified Eagle’s medium containing 2% FBS and 10 mM HEPES (pH 7.4). At 18 hpi, the cells were fixed by adding 100 µL of warm 1% paraformaldehyde (PFA; Electron Microscopy Science) in PBS and incubating for 1 hour. Following five washes with PBS, the cells were permeabilized using 0.1% saponin in PBS with 0.1% BSA. For immunostaining, CHIKV-, SFV-, ONNV-, and MAYV-infected cells were incubated with a rabbit pAb to E2/E1(50), RRV-infected cells with a rabbit polyclonal antibody against SFV capsid (51), and SINV-infected cells with monoclonal antibodies R2 and R6 targeting SINV E1 and E2 (52). Subsequently, cells were treated with horseradish peroxidase-conjugated goat anti-mouse or anti-rabbit IgG. Foci were visualized using TrueBlue Peroxidase substrate (#5510-0030, Seracare) and quantified with an ImmunoSpot S6 Macroanalyzer (Cellular Technologies).

### Dose-response antiviral assays

For CHIKV-nLuc–based dose-response assays, NHDF cells were seeded at 10,000 cells per well in 24-well plates and incubated for 24 hours under standard culture conditions. Cells were subsequently inoculated with the indicated virus at MOI = 0.1 FFU per cell using Med-A medium for 1 hour. Following infection, cells were washed three times with complete growth medium to eliminate unbound virus, and then treated with three-fold serial dilutions of compounds. Cells were lysed at 24 hpi, and nanoluciferase activity was measured using the Nano-Glo Dual-Luciferase Reporter Assay System (Promega) on a PerkinElmer Victor X5 multilabel plate reader. Dose-response data were analyzed by non-linear regression with a variable slope model to determine EC_50_ using GraphPad Prism 10. For virus yield–based dose-response analyses, U-2 OS cells were seeded at 75,000 cells per well in 24-well plates, inoculated with the indicated alphaviruses and treated with compounds as described above. At 24 hpi, supernatants were collected, clarified by centrifugation, and viral titers were determined using FFA. DMSO-treated cells were included as negative controls and nanoluciferase signals were normalized to the respective DMSO control values. Dose-response data were analyzed by normalized non-linear regression with a variable slope model to determine EC_50_ using GraphPad Prism 10.

### Selection for viral resistance

U-2 OS cells were seeded at a density of 200,000 cells per well in 6-well plates, cultured for approximately 24 hours, and then inoculated with CHIKV primary stock at MOI = 1 FFU/cell. After 2 hours, media containing compound or DMSO were added to three wells. At 16 hpi, supernatants were collected, clarified by centrifugation, frozen and viral titers were determined using FFA. The concentration of compounds in each passage is shown in Fig.6B. Viruses from P6 were plaque-purified, amplified in culture, and used for RNA isolation. Viral RNAs were reverse-transcribed and subjected to whole genome sequencing as below.

### Whole genome sequencing and data analysis

Viral whole-genome sequencing of CHIKV samples was performed using a metagenomic next-generation sequencing approach, as previously described (53). Libraries were sequenced on NextSeq 2000 with 2x150bp read format. Raw reads were trimmed and quality filtered with fastp (v0.23.4) (54) (--cut_mean_quality 20 --cut_front -- cut_tail --length_required 20 --low_complexity_filter --trim_poly_g --trim_poly_x). Filtered reads were then used for variant calling using the RAVA workflow (default parameters) and the CHIKV 181-Clone 25 (MK028839.1) as a reference (https://github.com/greninger-lab/RAVA_Pipeline/tree/2025-08-14_AECM_MK-PY_C3-4_CHIKV_publication) (55, 56).

### Cytotoxicity assay

NHDF cells were seeded at 10,000 cells per well in 24-well plates and cultured for 24 hours. Cells were then treated with three-fold serial dilutions of compounds starting at 500 µM and incubated for 24 hours. DMSO was included as a negative control for normalization. Following treatment, cells were incubated with PrestoBlue reagent (ThermoFisher Scientific) for 1 hour at 37°C, and fluorescence was measured using a PerkinElmer Victor X5 multilabel plate reader. The CC50 values were determined by normalized non-linear regression with a variable slope model in Prism (GraphPad).

### Trans-replication assays

U-2 OS cells were seeded at 75,000 cells per well in 24-well plates, and cultured for 24 hours. Cells were then co-transfected with 250 ng of plasmid encoding template RNA and 250 ng of plasmid encoding P1234 WT (57, 58), using Lipofectamine LTX with PLUS reagent (ThermoFisher Scientific) following the manufacturer’s instructions. After 4 hours of post-transfection, medium containing 100 µM of compounds or DMSO was added to the cells. Cells were then incubated for 20 hours, lysed, and Gluc activity was quantified using the Dual-Luciferase Reporter Assay System (Promega). Luciferase activities were normalized to those of the DMSO-treated controls.

### In silico Docking

Docking studies were performed with MOE 2024.06, using the Amber10 force field. A homology model of CHIKV nsP4 based on the coordinates reported for SINV nsP4 (PDB 7VW5) was used for docking studies. After protonation and energy minimization, an induced-fit protocol was used to dock GAP-1146924 and GAP-1173149 into the CHIKV nsP4 structure based on resistance data information. For GAP-1146924, residue 542 was selected to identify the target site for docking. For GAP-1173149, residue A350 was selected as the target site for docking.

### Synthesis of 2-(2,5-dioxoimidazolidin-1-yl)-N-(1,4,5,6,7,8-hexahydrocyclohepta[d] pyrazol-3-ylmethyl)acetamide (GAP-1173149)

2-(2,5-dioxoimidazolidin-1-yl)acetic acid was dissolved in 30 mL DMF and to the reaction added 5.06 g (13.31 mmol 1.1 eq) HATU and the reaction mixture was brought to 0°C. Then the amine 1,4,5,6,7,8-hexahydrocyclohepta[d]pyrazol-3-ylmethanamine (2g, 12.1 mmol) and 2.11 mL (12 mmol, 1 eq) DIPEA was added and stirred while letting the mixture come to room temperature for 6 hours. After checking the reaction completion by thin layer chromatography and staining with PMA, and LC-MS, the reaction was terminated by the addition of a few drops of HCl solution, and extracted 2 times with cold water and ethyl acetate. The organic layers were collected, dried with magnesium sulfate, filtered and evaporated. The residue was purified using column chromatography with DCM and increasing ratio of DCM:MeOH:Ammonnia (9:1:0.2) mixture and then with reverse phase using C18 reverse phase column water/acetonitrile system to yield 1.3 g 2-(2,5-dioxoimidazolidin-1-yl)-N-(1,4,5,6,7,8-hexahydrocyclohepta[d]pyrazol-3-ylmethyl)acetamide (GAP-1173149) with 35% yield.

### Synthesis of 2-(4-fluorophenyl)-N-[(2-methylpyrazol-3-yl)methyl]ethanamine Intermediate

2-(4-fluorophenyl)ethanamine (1g, 7.19 mmol) was dissolved in anhydrous DCM. 2-methylpyrazole-3-carbaldehyde (791mg, 7.19 mmol, 1 eq) and titanium (IV) isopropoxide (2.53 mL, 8.62 mmol, 1.2 eq) were added to the reaction and stirred under argon atmosphere for 6 hours. After this time, 3.81 g (17.96 mmol, 2.5 eq) sodium triacetoxyborohydride and catalytic amount acetic acid were added and stirred overnight. The reaction was then brought to pH 10 with NaOH solution. More DCM and water were added. Phases separated, organic layers collected, dried with magnesium sulfate, filtered and evaporated under vacuum. Product were purified using flash chromatography with hexanes and increasing ratio of ethyl acetate to obtain 1.24 g 2-(4-fluorophenyl)-N-[(2-methylpyrazol-3-yl)methyl]ethanamine with 74% yield.

### Synthesis of N-[2-(4-fluorophenyl)ethyl]-1,5-dimethyl-N-[(2-methylpyrazol-3-yl)methyl]pyrazole-4-sulfonamide (GAP-1146924)

2-(4-fluorophenyl)-N-[(2-methylpyrazol-3-yl)methyl]ethanamine (1.24 g, 5.32 mmol) was dissolved in 20 mL anhydrous THF and N,N-DIPEA (1.85 mL, 10.63 mmol, 2eq) added. Reaction was brought to 0°C, and then 1.14 g (5.85 mmol, 1.1 eq) 1,5-dimethylpyrazole-4-sulfonyl chloride was added. Reaction was let to warm up to room temperature and stirred for 3h in total. After the reaction was complete, ethyl acetate and water was added for extraction. Organic layers were collected, dried using magnesium sulfate, filtered and evaporated under vacuum. Product were purified using flash chromatography with hexanes and increasing ratio of ethyl acetate to obtain 1.2 g N-[2-(4-fluorophenyl)ethyl]-1,5-dimethyl-N-[(2-methylpyrazol-3-yl)methyl]pyrazole-4-sulfonamide (GAP-1146924) with 60% yield.

**NMR analysis of GAP-1173149.** ^1^H NMR (400 MHz, MeOD) δ 4.34 (s, 2H), 4.13 (s, 2H), 4.01 (s, 2H), 2.74 - 2.67 (m, 2H), 2.59 - 2.49 (m, 2H), 1.85 (qd, *J* = 6.1, 3.3 Hz, 2H), 1.71 - 1.59 (m, 4H). (NH protons exchanged with MeOD). ^13^C NMR (151 MHz, MeOD) δ 172.48, 167.28, 158.10, 147.99, 142.30, 117.87, 46.20, 39.88, 34.11, 31.90, 28.71, 27.31, 27.28, 23.62.

**NMR analysis of 2-(4-fluorophenyl)-N-[(2-methylpyrazol-3-yl)methyl]ethanamine Intermediate.** ^1^H NMR (400 MHz, CDCl3) δ 7.39 (d, J = 1.8 Hz, 1H), 7.21 - 7.12 (m, 2H), 7.06 - 6.92 (m, 2H), 6.13 (d, J = 1.8 Hz, 1H), 3.84 (s, 3H), 3.82 (s, 2H), 2.90 (t, J = 1.1 Hz, 2H), 2.80 (t, J = 6.9 Hz, 2H), 1.82 (s, 1H). ^19^F NMR (376 MHz, CDCl3) δ -116.97.

**NMR analysis of GAP-1146924.** ^1^H NMR (400 MHz, DMSO) δ 7.81 (s, 1H), 7.37 (d, *J* = 1.8 Hz, 1H), 7.10 - 6.95 (m, 4H), 6.28 (d, *J* = 1.8 Hz, 1H), 4.35 (s, 2H), 3.79 (s, 3H), 3.75 (s, 3H), 3.20 - 3.10 (m, 2H), 2.52 - 2.49 (m, 2H), 2.44 (s, 3H). ^19^F NMR (376 MHz, DMSO) δ -116.80. ^13^C NMR (101 MHz, DMSO) δ 161.31 (d, *J* = 241.7 Hz, C-F)), 141.01, 138.60, 137.91, 137.04, 134.85 (d, *J* = 3.1 Hz, C-C-C-C-F), 130.72 (d, *J* = 8.0 Hz, C-C-C-F), 116.61, 115.49 (d, *J* = 21.1 Hz, C-C-F) 107.73, 49.51, 43.32, 37.21, 36.72, 34.18, 10.29.

### Statistics and Reproducibility

CBIS, GraphPad Prism (10.3.0), and Excel (16.92) software packages were used for data analysis. Statistical significance was assessed using ordinary one-way ANOVA with multiple comparisons test in GraphPad Prism. For antiviral potency and cytotoxicity measurements, effective concentrations were determined from dose-response data sets through 4-parameter variable slope regression modeling. Biological repeats indicate measurements taken from distinct samples.

## Data availability statement

CHIKV whole genome sequencing data of the compound-treated and DMSO treated passages are available in NCBI BioProject PRJNA1306170. HTS and dose response raw data are available from the corresponding authors upon request. Numerical data for all biological experiments associated with the individual figures are available from the corresponding authors upon request. However, no chemical structure information concerning the composition of screening libraries and unsuccessful hit candidates will be provided.

## Code availability statement

Homology modeling software is available from the AlphaFold modeling server at https://alphafoldserver.com/. All other software solutions employed are commercially available.

## Acknowledgements

Chemdraw was employed for drawing, displaying, and characterizing chemical structures, substructures, and reactions (PerkinElmer21.0.0). This work was supported by U19 grant AI171403 (to RKP and GR Painter) under Project 2 (to MK and AM) and Scientific Cores D (to ALG) and F (to RMC). The content of this paper is solely the responsibility of the authors and does not necessarily represent the official views of the NIH, NIAID, or NCI. The funders had no role in study design, data collection and interpretation, or the decision to submit the work for publication.

## References

1. Kuhn R. 2021. Togaviridae: The Viruses and Their Replication., p 170-93. In Howley PM KD (ed), Fields Virology: Emerging Viruses, 7th ed, vol 1. Lippincott Williams & Wilkins, Philadelphia, PA.

2. Strauss JH, Strauss EG. 1994. The alphaviruses: gene expression, replication, and evolution. Microbiol Rev 58:491–562.

3. de Souza WM, Lecuit M, Weaver SC. 2025. Chikungunya virus and other emerging arthritogenic alphaviruses. Nat Rev Microbiol 23:585–601.

4. Levi LI, Vignuzzi M. 2019. Arthritogenic Alphaviruses: A Worldwide Emerging Threat? Microorganisms 7.

5. Silva LA, Dermody TS. 2017. Chikungunya virus: epidemiology, replication, disease mechanisms, and prospective intervention strategies. J Clin Invest 127:737–749.

6. de Souza WM, Fumagalli MJ, de Lima STS, Parise PL, Carvalho DCM, Hernandez C, de Jesus R, Delafiori J, Candido DS, Carregari VC, Muraro SP, Souza GF, Simoes Mello LM, Claro IM, Diaz Y, Kato RB, Trentin LN, Costa CHS, Maximo A, Cavalcante KF, Fiuza TS, Viana VAF, Melo MEL, Ferraz CPM, Silva DB, Duarte LMF, Barbosa PP, Amorim MR, Judice CC, Toledo-Teixeira DA, Ramundo MS, Aguilar PV, Araujo ELL, Costa FTM, Cerqueira-Silva T, Khouri R, Boaventura VS, Figueiredo LTM, Fang R, Moreno B, Lopez-Verges S, Mello LP, Skaf MS, Catharino RR, Granja F, Martins-de-Souza D, Plante JA, Plante KS, Sabino EC, Diamond MS, et al. 2024. Pathophysiology of chikungunya virus infection associated with fatal outcomes. Cell Host Microbe 32:606–622 e8.

7. Cerqueira-Silva T, Pescarini JM, Cardim LL, Leyrat C, Whitaker H, Antunes de Brito CA, Brickley EB, Barral-Netto M, Barreto ML, Teixeira MG, Boaventura VS, Paixao ES. 2024. Risk of death following chikungunya virus disease in the 100 Million Brazilian Cohort, 2015-18: a matched cohort study and self-controlled case series. Lancet Infect Dis 24:504–513.

8. Zacks MA, Paessler S. 2010. Encephalitic alphaviruses. Vet Microbiol 140:281–6.

9. Lindsey NP, Martin SW, Staples JE, Fischer M. 2020. Notes from the Field: Multistate Outbreak of Eastern Equine Encephalitis Virus - United States, 2019. MMWR Morb Mortal Wkly Rep 69:50–51.

10. Morrison TE. 2014. Reemergence of chikungunya virus. J Virol 88:11644–7.

11. de Souza WM, Ribeiro GS, de Lima STS, de Jesus R, Moreira FRR, Whittaker C, Sallum MAM, Carrington CVF, Sabino EC, Kitron U, Faria NR, Weaver SC. 2024. Chikungunya: a decade of burden in the Americas. Lancet Reg Health Am 30:100673.

12. McCarthy MK, Davenport BJJ, Morrison TE. 2022. Chronic Chikungunya Virus Disease. Curr Top Microbiol Immunol 435:55–80.

13. Tsetsarkin KA, Chen R, Weaver SC. 2016. Interspecies transmission and chikungunya virus emergence. Curr Opin Virol 16:143–150.

14. Tsetsarkin KA, Vanlandingham DL, McGee CE, Higgs S. 2007. A single mutation in chikungunya virus affects vector specificity and epidemic potential. PLoS Pathog 3:e201.

15. Codeco CT, Oliveira SS, Ferreira DAC, Riback TIS, Bastos LS, Lana RM, Almeida IF, Godinho VB, Cruz OG, Coelho FC. 2022. Fast expansion of dengue in Brazil. Lancet Reg Health Am 12:100274.

16. Ryan SJ, Carlson CJ, Mordecai EA, Johnson LR. 2019. Global expansion and redistribution of Aedes-borne virus transmission risk with climate change. PLoS Negl Trop Dis 13:e0007213.

17. Fournier L, Durand GA, Cochet A, Brottet E, Fiet C, Mano Q, Krug C, Verdurme L, Blanchot T, Fournier R, Paty MC, Grard G, Franke F, Calba C. 2025. Multiple early local transmissions of chikungunya virus, Mainland France, from May 2025. Euro Surveill 30.

18. Wang J, Zhang L. 2025. The chikungunya virus outbreak in Foshan, China: A rising public health threat in tropical and subtropical regions. J Infect 91:106591.

19. Burt FJ, Rolph MS, Rulli NE, Mahalingam S, Heise MT. 2012. Chikungunya: a re-emerging virus. Lancet 379:662–71.

20. Luo D, Tan YB, Law MCY, Jin J. 2025. A Structural Perspective on the Alphavirus Life Cycle. Annu Rev Virol doi:10.1146/annurev-virology-093022-010359.

21. Ahola T, McInerney G, Merits A. 2021. Alphavirus RNA replication in vertebrate cells. Adv Virus Res 111:111–156.

22. Law YS, Utt A, Tan YB, Zheng J, Wang S, Chen MW, Griffin PR, Merits A, Luo D. 2019. Structural insights into RNA recognition by the Chikungunya virus nsP2 helicase. Proc Natl Acad Sci U S A 116:9558–9567.

23. Ghoshal A, Asressu KH, Hossain MA, Brown PJ, Nandakumar M, Vala A, Merten EM, Sears JD, Law I, Burdick JE, Morales NL, Perveen S, Pearce KH, Popov KI, Moorman NJ, Heise MT, Willson TM. 2024. Structure Activity of beta-Amidomethyl Vinyl Sulfones as Covalent Inhibitors of Chikungunya nsP2 Cysteine Protease with Antialphavirus Activity. J Med Chem 67:16505–16532.

24. Metibemu DS, Adeyinka OS, Falode J, Hampton T, Crown O, Ojobor JC, Narayanan A, Julander J, Ogungbe IV. 2024. Inhibitor of the non-structural protein 2 protease shows promising efficacy in mouse models of chikungunya. Eur J Med Chem 278:116808.

25. Abu Bakar F, Ng LFP. 2018. Nonstructural Proteins of Alphavirus-Potential Targets for Drug Development. Viruses 10.

26. Delang L, Segura Guerrero N, Tas A, Querat G, Pastorino B, Froeyen M, Dallmeier K, Jochmans D, Herdewijn P, Bello F, Snijder EJ, de Lamballerie X, Martina B, Neyts J, van Hemert MJ, Leyssen P. 2014. Mutations in the chikungunya virus non-structural proteins cause resistance to favipiravir (T-705), a broad-spectrum antiviral. J Antimicrob Chemother 69:2770–84.

27. Ehteshami M, Tao S, Zandi K, Hsiao HM, Jiang Y, Hammond E, Amblard F, Russell OO, Merits A, Schinazi RF. 2017. Characterization of beta-d-N(4)-Hydroxycytidine as a Novel Inhibitor of Chikungunya Virus. Antimicrob Agents Chemother 61.

28. Ferreira AC, Reis PA, de Freitas CS, Sacramento CQ, Villas Boas Hoelz L, Bastos MM, Mattos M, Rocha N, Gomes de Azevedo Quintanilha I, da Silva Gouveia Pedrosa C, Rocha Quintino Souza L, Correia Loiola E, Trindade P, Rangel Vieira Y, Barbosa-Lima G, de Castro Faria Neto HC, Boechat N, Rehen SK, Bruning K, Bozza FA, Bozza PT, Souza TML. 2019. Beyond Members of the Flaviviridae Family, Sofosbuvir Also Inhibits Chikungunya Virus Replication. Antimicrob Agents Chemother 63.

29. Yin P, May NA, Lello LS, Fayed A, Parks MG, Drobish AM, Wang S, Andrews M, Sticher Z, Kolykhalov AA, Natchus MG, Painter GR, Merits A, Kielian M, Morrison TE. 2024. 4’-Fluorouridine inhibits alphavirus replication and infection in vitro and in vivo. mBio 15:e0042024.

30. Yin P, Sobolik EB, May NA, Wang S, Fayed A, Vyshenska D, Drobish AM, Parks MG, Lello LS, Merits A, Morrison TE, Greninger AL, Kielian M. 2025. Mutations in chikungunya virus nsP4 decrease viral fitness and sensitivity to the broad-spectrum antiviral 4’-Fluorouridine. PLoS Pathog 21:e1012859.

31. Abdelnabi R, Kovacikova K, Moesslacher J, Donckers K, Battisti V, Leyssen P, Langer T, Puerstinger G, Querat G, Li C, Decroly E, Tas A, Marchand A, Chaltin P, Coutard B, van Hemert M, Neyts J, Delang L. 2020. Novel Class of Chikungunya Virus Small Molecule Inhibitors That Targets the Viral Capping Machinery. Antimicrob Agents Chemother 64.

32. Battisti V, Moesslacher J, Abdelnabi R, Leyssen P, Rosales Rosas AL, Langendries L, Aufy M, Studenik C, Kratz JM, Rollinger JM, Puerstinger G, Neyts J, Delang L, Urban E, Langer T. 2024. Design, synthesis, and lead optimization of piperazinyl-pyrimidine analogues as potent small molecules targeting the viral capping machinery of Chikungunya virus. Eur J Med Chem 264:116010.

33. Lucas CJ, Davenport BJ, Carpentier KS, Tinega AN, Morrison TE. 2022. Two Conserved Phenylalanine Residues in the E1 Fusion Loop of Alphaviruses Are Essential for Viral Infectivity. J Virol 96:e0006422.

34. Couderc T, Lecuit M. 2009. Focus on Chikungunya pathophysiology in human and animal models. Microbes Infect 11:1197–205.

35. Abramson J, Adler J, Dunger J, Evans R, Green T, Pritzel A, Ronneberger O, Willmore L, Ballard AJ, Bambrick J, Bodenstein SW, Evans DA, Hung CC, O’Neill M, Reiman D, Tunyasuvunakool K, Wu Z, Zemgulyte A, Arvaniti E, Beattie C, Bertolli O, Bridgland A, Cherepanov A, Congreve M, Cowen-Rivers AI, Cowie A, Figurnov M, Fuchs FB, Gladman H, Jain R, Khan YA, Low CMR, Perlin K, Potapenko A, Savy P, Singh S, Stecula A, Thillaisundaram A, Tong C, Yakneen S, Zhong ED, Zielinski M, Zidek A, Bapst V, Kohli P, Jaderberg M, Hassabis D, Jumper JM. 2024. Accurate structure prediction of biomolecular interactions with AlphaFold 3. Nature 630:493–500.

36. Gane EJ, Stedman CA, Hyland RH, Ding X, Svarovskaia E, Symonds WT, Hindes RG, Berrey MM. 2013. Nucleotide polymerase inhibitor sofosbuvir plus ribavirin for hepatitis C. N Engl J Med 368:34–44.

37. Sulkowski MS, Gardiner DF, Rodriguez-Torres M, Reddy KR, Hassanein T, Jacobson I, Lawitz E, Lok AS, Hinestrosa F, Thuluvath PJ, Schwartz H, Nelson DR, Everson GT, Eley T, Wind-Rotolo M, Huang SP, Gao M, Hernandez D, McPhee F, Sherman D, Hindes R, Symonds W, Pasquinelli C, Grasela DM, Group AIS. 2014. Daclatasvir plus sofosbuvir for previously treated or untreated chronic HCV infection. N Engl J Med 370:211–21.

38. Koff RS. 2014. Review article: the efficacy and safety of sofosbuvir, a novel, oral nucleotide NS5B polymerase inhibitor, in the treatment of chronic hepatitis C virus infection. Aliment Pharmacol Ther 39:478–87.

39. Lupina K, Nowak K, Lorek D, Nowak A, Romac A, Glowacka E, Janczura J. 2025. Pharmacological advances in HIV treatment: from ART to long-acting injectable therapies. Arch Virol 170:195.

40. Painter GR, Natchus MG, Cohen O, Holman W, Painter WP. 2021. Developing a direct acting, orally available antiviral agent in a pandemic: the evolution of molnupiravir as a potential treatment for COVID-19. Curr Opin Virol 50:17–22.

41. Li G, Wang Y, De Clercq E. 2022. Approved HIV reverse transcriptase inhibitors in the past decade. Acta Pharm Sin B 12:1567–1590.

42. Cox RM, Sourimant J, Toots M, Yoon JJ, Ikegame S, Govindarajan M, Watkinson RE, Thibault P, Makhsous N, Lin MJ, Marengo JR, Sticher Z, Kolykhalov AA, Natchus MG, Greninger AL, Lee B, Plemper RK. 2020. Orally efficacious broad-spectrum allosteric inhibitor of paramyxovirus polymerase. Nat Microbiol 5:1232–1246.

43. Yan D, Weisshaar M, Lamb K, Chung HK, Lin MZ, Plemper RK. 2015. Replication-Competent Influenza Virus and Respiratory Syncytial Virus Luciferase Reporter Strains Engineered for Co-Infections Identify Antiviral Compounds in Combination Screens. Biochemistry 54:5589–604.

44. Ashbrook AW, Burrack KS, Silva LA, Montgomery SA, Heise MT, Morrison TE, Dermody TS. 2014. Residue 82 of the Chikungunya virus E2 attachment protein modulates viral dissemination and arthritis in mice. J Virol 88:12180–92.

45. Liljeström P, Lusa S, Huylebroeck D, Garoff H. 1991. In vitro mutagenesis of a full-length cDNA clone of Semliki Forest virus: the small 6,000-molecular-weight membrane protein modulates virus release. J Virol 65:4107–13.

46. Hardwick JM, Levine B. 2000. Sindbis virus vector system for functional analysis of apoptosis regulators. Methods Enzymol 322:492–508.

47. Cox RM, Toots M, Yoon JJ, Sourimant J, Ludeke B, Fearns R, Bourque E, Patti J, Lee E, Vernachio J, Plemper RK. 2018. Development of an allosteric inhibitor class blocking RNA elongation by the respiratory syncytial virus polymerase complex. J Biol Chem 293:16761–16777.

48. Baell JB, Nissink JWM. 2018. Seven Year Itch: Pan-Assay Interference Compounds (PAINS) in 2017-Utility and Limitations. ACS Chem Biol 13:36–44.

49. Yang JJ, Ursu O, Lipinski CA, Sklar LA, Oprea TI, Bologa CG. 2016. Badapple: promiscuity patterns from noisy evidence. J Cheminform 8:29.

50. Kielian M, Jungerwirth S, Sayad KU, DeCandido S. 1990. Biosynthesis, maturation, and acid activation of the Semliki Forest virus fusion protein. J Virol 64:4614–24.

51. Brown RS, Anastasakis DG, Hafner M, Kielian M. 2020. Multiple capsid protein binding sites mediate selective packaging of the alphavirus genomic RNA. Nat Commun 11:4693.

52. Meyer WJ, Gidwitz S, Ayers VK, Schoepp RJ, Johnston RE. 1992. Conformational alteration of Sindbis virion glycoproteins induced by heat, reducing agents, or low pH. J Virol 66:3504–13.

53. Greninger AL, Zerr DM, Qin X, Adler AL, Sampoleo R, Kuypers JM, Englund JA, Jerome KR. 2017. Rapid Metagenomic Next-Generation Sequencing during an Investigation of Hospital-Acquired Human Parainfluenza Virus 3 Infections. J Clin Microbiol 55:177–182.

54. Chen S, Zhou Y, Chen Y, Gu J. 2018. fastp: an ultra-fast all-in-one FASTQ preprocessor. Bioinformatics 34:i884–i890.

55. Di Tommaso P, Chatzou M, Floden EW, Barja PP, Palumbo E, Notredame C. 2017. Nextflow enables reproducible computational workflows. Nat Biotechnol 35:316–319.

56. Lin MJ, Shean RC, Makhsous N, Greninger AL. 2019. LAVA: a streamlined visualization tool for longitudinal analysis of viral alleles. bioRxiv doi:10.1101/2019.12.17.879320:2019.12.17.879320.

57. Lello LS, Miilimäe A, Cherkashchenko L, Omler A, Skilton R, Ireland R, Ulaeto D, Merits A. 2023. Activity, Template Preference, and Compatibility of Components of RNA Replicase of Eastern Equine Encephalitis Virus. J Virol 97:e0136822.

58. Utt A, Rausalu K, Jakobson M, Männik A, Alphey L, Fragkoudis R, Merits A. 2019. Design and Use of Chikungunya Virus Replication Templates Utilizing Mammalian and Mosquito RNA Polymerase I-Mediated Transcription. J Virol 93.

